# ETAPOD: A forecast model for prediction of black pod disease Outbreak in Nigeria

**DOI:** 10.1101/488452

**Authors:** Peter M. Etaware, Adegboyega R. Adedeji, Oyedeji I. Osowole, Adegboyega C. Odebode

## Abstract

The misuse of toxic fungicides by indigenous cocoa farmers in Nigeria stem from their inability to predict the time for black pod disease (BPD) outbreak. Prediction of possible time for BPD outbreak will provide spotlight on areas under massive BPD invasion, minimise fungicide misuse and increase control accuracy. The Multiple Regression Model (MRM): Y=α+β_1_X_1_+β_2_X_2_+…+β_n_X_n_ where Y is Nx1 matrix of response variable, X_1_,X_2_,…X_n_ are NxK matrices of regressors, and β_1_,β_2_,…β_n_ regression coefficients was used in model development. Eight models (MRM_1_-MRM_8_) were fitted from real life BPD data. The performances of the models were ascertained using SER, RMSE_pred_ and R-Sq_Adj_. Prediction(s) made by the best fitted model was compared to real life observations (Monthly BPD Occurrence (MBO), Total Annual Occurrence (TAO), and Average Annual Occurrence (AAO), respectively). The preferred model was MRM_5_ (ETAPOD) followed by MRM_4_, MRM_1_, MRM_2_, and MRM_3_ in terms of SER (0.22, 0.39, 0.45, 0.45 and 0.45), RMSE_pred_.(0.30, 039, 0.46, 0.46 and 0.46) and R-Sq_Ad_j.(0.67, 0.49, 0.32, 0.32, and 0.31), respectively. Predictions on BPD outbreak made by ETAPOD showed that MBO, TAO and AAO for some selected stations i.e. Ọwenà and Wáàsimi were 9.05, 72.3 and 6.0% compared with observed BPD values of 9.5, 70.0, and 5.8%, respectively. Adaàgbà, Iyánfoworogi, and Owódé-Igàngán had 9.43, 77.8, and 6.5% as their predicted BPD values compared with the observed values of 9.0, 53.5, and 4.46%, respectively. ETAPOD performed better than other models and its predicted values were within the range of real life occurrence.

## Introduction

Black pod disease of *Theobroma cacao* Linn. (Cocoa) is a major challenge to farmers worldwide. The disease is more established in West Africa than in any other parts of the world. As a result of the disease, most indigenous cocoa farmers seemingly have no enthusiasm in establishing new farms in areas where black pod disease outbreak is extreme [1] and cocoa farmlands are rapidly being abandoned [2]. Adegbola [3] in his review of Africa estimated the average occurrence of the disease as 40% in several parts of West Africa and up to 90% in certain places in Nigeria [4].

Global climate change is one of the major factors responsible for the irregular occurrence of this disease worldwide due to its influence on the survival and proliferation of the pathogen and the predisposition of unripe and ripe cocoa pods to fungal attack. The irregular rainfall pattern and inconsistent mode of black pod disease occurrence in Nigeria makes it nearly impossible to control it effectively. The success rate achieved by both biological and chemical control measures is fast declining due to high level of adaptation of the pathogen to harsh conditions, climate change and an increase in vulnerability of cocoa plants. Hence, an urgent need for modern approach in the control of black pod disease in West Africa is imminent.

Plant disease forecasting (which is a modern day plant disease control strategy) advocates the use of plethora management techniques directed by a rational system for predicting the risk of disease outbreaks such that farmers will be duly informed and maximally equipped either to avert or tackle the impending epidemics. This research work seeks to develop a forecast system for black pod disease prediction in order to provide useful and timely information on black pod disease outbreak. This will minimize fungicide misuse, increase cocoa productivity, and reduce the risk of chemical poisoning. Unless concerted efforts are made to effectively manage the disease [5], black pod disease epidemics will greatly reduce cocoa production in Nigeria and around the world [1].

## Materials and Methods

### Research location

Twelve study locations within Southwest, Nigeria were selected i.e. Ọwenà (Two study locations) and Wáàsimi (Ondo East Local Government Area, Ondo State), Adaàgbà and Iyánfọwọrọgi (Ife South L.G.A, Osun State), two study locations in Owódé-Igàngán (Àtàkúnmòsà East L.G.A., Osun State), two study locations in Ọbáfẹmi-Owódé L.G.A., Ogun State, Mòyè village Ọnà-Arà LGA, Dáagi-Lógbà (Atérè Village) in Ọmi-Adió area of Ìddó LGA, and Olórò Village (also known as Ọlọrunda village) in Àkànràn, Ọnà-Arà LGA of Oyo State, with the exception of Ekiti and Lagos States (Table 1).

**Table 1:** The State and Local Government Area of the sample stations in Southwest, Nigeria

| Location of Cocoa Farms | No. of Sample Stations | Local Govt. Area | State |
| --- | --- | --- | --- |
| Ọbáfẹmi-Owódé | 2 | Abeokuta | Ogun |
| Adaàgbà | 1 | Ife South | Osun |
| Owódé-Igàngán | 2 | Àtákúnmòsà East | Osun |
| Iyánfọwọrọgi | 1 | Ife South | Osun |
| Ọwenà | 2 | Ondo East | Ondo |
| Wáàsìmi | 1 | Ondo East | Ondo |
| Mòyè village | 1 | Ọnà-Arà | Oyo |
| Dáagi-Lógbà | 1 | Ìddó | Oyo |
| Olórò village | 1 | Ọnà-Arà | Oyo |

**Data Source:** BPD assessment (2015/2016) cocoa production season in Southwest, Nigeria

### The co-ordinates of the research locations

The co-ordinates of the study areas were determined using the blackberry mobile Global Positioning System (GPS) device (version 6.0) and a mobile satellite GPS receiver model GARMIN Etrex 10 obtained from the Department of Botany, Faculty of Science, University of Ibadan, Ibadan, Nigeria. The farm size was measured using a surveyor’s measuring tape (100ft by 30m) Lufkin FM100CME 2-Sided, Metric/English 13mm ½ inch x (Fig 1).

**Fig 1:**
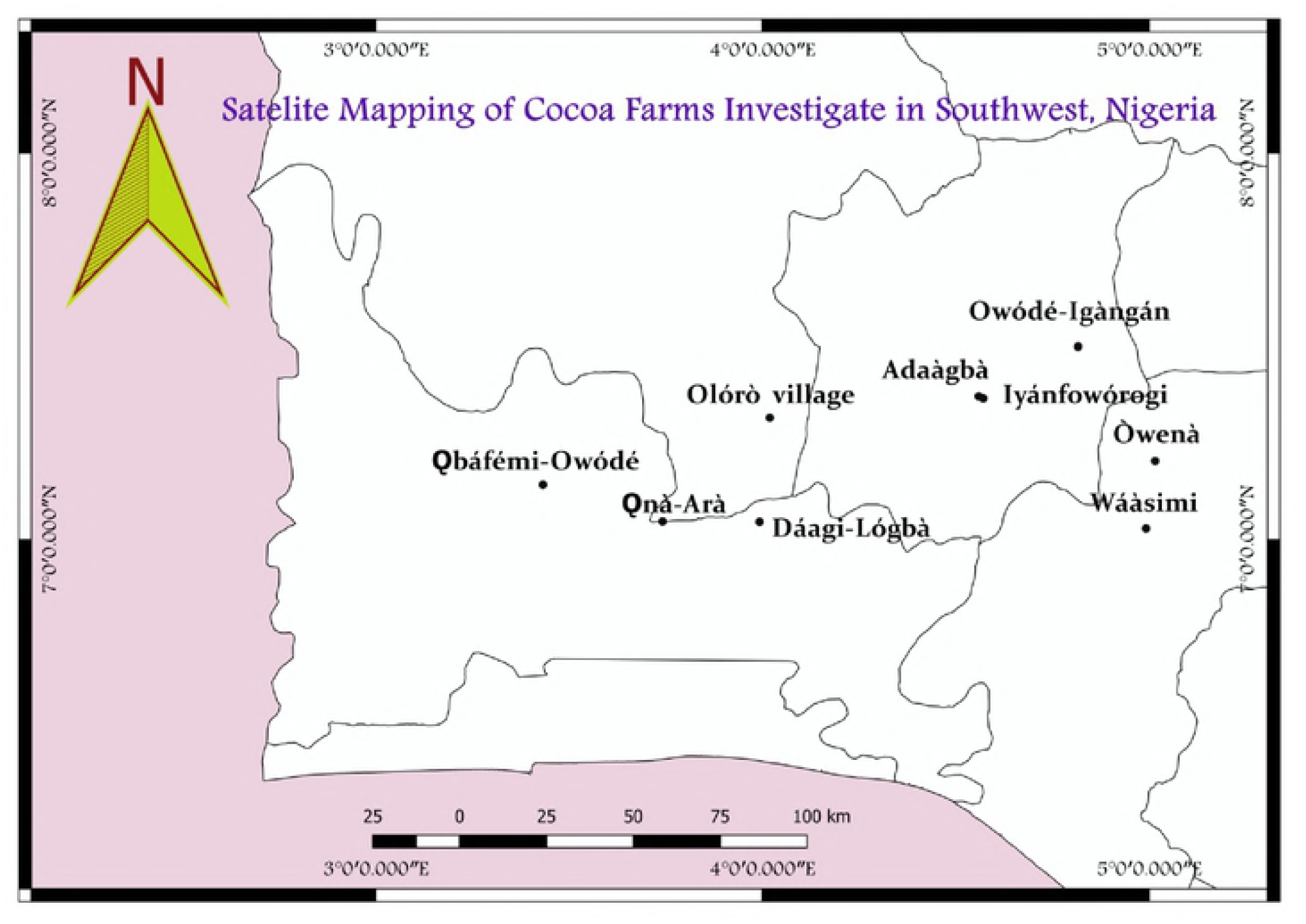
Coordinates of cocoa farm site where primary data were collected for this research

### Black pod disease data

Documented reports of black pod disease outbreaks within Southwestern Nigeria was obtained from Cocoa Research Institute of Nigeria (CRIN), Ìdí-Ayunrẹ, Ibadan, Oyo State, Nigeria and the report of Lawal and Emaku [6]. The total data collected spanned from 1985 to 2014. These served as the secondary data while the primary data were directly collected in the field during disease assessment (2015/2016).

### Meteorological data

Weather data from 1985 to 2014 were also collected from the National Bureau of Statistics (NBS) Ibadan, Oyo State, the Meteorological Station of Cocoa Research Institute of Nigeria (CRIN), Ìdí-Ayunrẹ, Ibadan, Oyo State, Nigerian Meteorological Station (Nimet), Nigerian Institute for Oil palm Research (NIFOR), Benin City, Edo State, Nigeria, and the Department of Geography, University of Ibadan, Ibadan, Oyo State, Nigeria. These were also classified as secondary data.

### Forecast model structuring

The secondary data (BPD outbreak and weather data) were used in the structuring of the proposed forecast model, while the primary data gotten directly from the field (2015/2016) was used to validate the predicted results for black pod disease outbreak by the developed model for the same period. The template for validation was stated in Table 2.

**Table 2:** Cross Validation (CV) format

**(© Etaware Peter Mudiaga)**
| **BPD Outbreak (Live)** | **BPD Outbreak (Predicted)** | **Inference** |
| --- | --- | --- |
| 100 Cocoa stands affected | 100% Disease Prediction | 100% Accuracy |
| 100 Cocoa stands affected | 90% Disease Prediction | 90% Accuracy |
| 100 Cocoa stands affected | 80% Disease Prediction | 80% Accuracy |
| 100 Cocoa stands affected | 70% Disease Prediction | 70% Accuracy |
| 100 Cocoa stands affected | 60% Disease Prediction | 60% Accuracy |
| 100 Cocoa stands affected | 50% Disease Prediction | 50% Accuracy |
| 100 Cocoa stands affected | 40% Disease Prediction | 40% Accuracy |
| 100 Cocoa stands affected | 30% Disease Prediction | 30% Accuracy |
| 100 Cocoa stands affected | 20% Disease Prediction | 20% Accuracy |
| 100 Cocoa stands affected | 10% Disease Prediction | 10% Accuracy |
| 100 Cocoa stands affected | 0% Disease Prediction | 0% Accuracy |

(© Etaware Peter Mudiaga)

### Data Analysis

The proposed forecast model(s) were templates of multiple regression equation(s) developed from the meteorological data and previous black pod disease records collected (Secondary data), designed using Minitab 16.0 software and programmed on Microsoft Excel Worksheet 2007 service pack for easy access. The validity of the developed models was tested using Pearson’s Product Moment of Correlation (PPM) to determine the Coefficient of Correlation (R-Sq), and the Adjusted Coefficient of Correlation of the Developed Models (R-Sq_Adj_). The Standard Error of Regression (SER) and Root Mean Square Error of Prediction (RMSE_pred_.) were also determined as a valid tool for black pod disease forecast model selection.

Predicted results of black pod disease outbreak were validated using the observations made on the field (Primary data) during the 2015/2016 black pod disease assessments in the study areas. The Error of in prediction was also determined using E=(Y-Ŷ)^2^. Qualitative data were represented as charts and graphs plotted using SPSS, version 20.0 for 32 bits resolution, while the analysis of variance was carried out using COSTAT 9.0 software. The homogeneity of means was determined using Duncan Multiple Range Test (DMRT).

## Results

The developed black pod disease function was modeled using simple mathematical rule

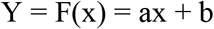

Where, a = Integer, x = independent variable, b = Constant, Y = Response variable and F = The Function of the variable x

Thus,

BPD Outbreak = F(Host x Pathogen x Environment) = a(Host x Pathogen x Environment) + b

Mathematically,

Recall, Y = F(x) = ax + b

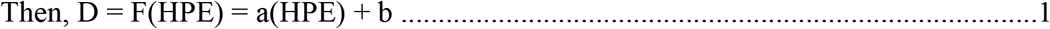

In any case the influence of man and vectors (Ants, Termites, and Rodents etc.) serve as constants in the equation because they influence the spread of black pod disease in the field, coupled with the timely combination of the key factors responsible for black pod disease development.

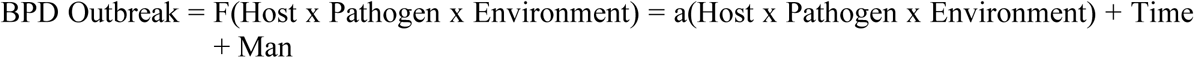

Mathematically,

Recall, Y = F(x) = ax + b

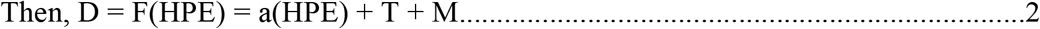

Where, D = BPD Outbreak, H = Host, P = Pathogen, T = Time, M = man, E = Environment, x = HPE, BPD = Black pod disease, and b = (T + M)

Therefore, if the disease equation is differentiated with respect to the timing of occurrence, then the equation below becomes a derivative of the first order differentiation for BPD outbreak.

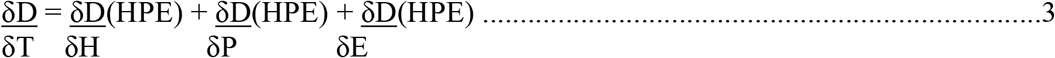

The forecast models were structured using the Multiple Regression Equation (MRM):

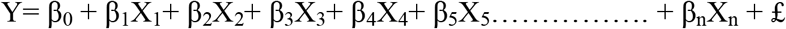

Since α = β_0_, 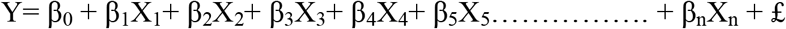

Where, Y = Response variable, X_1_, X_2_, X_3_, X_4_, X_5_,* X_n_ = Predictors, β_1_, β_2_, β_3_, β_4_, β_5_ β_n_ = The slopes, α = General constant and £ = The error factor for the predictors [7]

Therefore, the development of black pod disease forecast system for cocoa required an equation encompassing all the predictors necessary for the disease development. An example of such model was given thus:

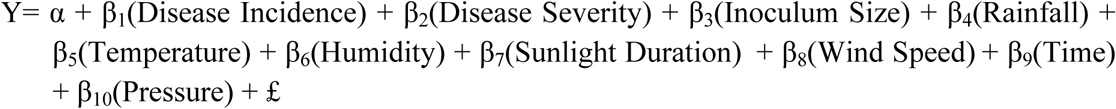

Or

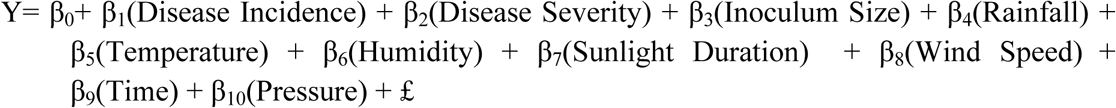

In any case the individual predictors were tested against the response variable to ascertain their role(s) in black pod disease outbreak. In some cases the relationship of a predictor to the response variable was in the reverse order, this was still acceptable. In a situation whereby a chosen predictor has no established relationship with the response variable, then that predictor was discarded (Fig 2).

Rainfall and average relative humidity had a positive correlation with BPD outbreak (r = 0.445 and 0.477, and r^2^ = 0.105 and 0.295, respectively) as shown in Figs 3 and 5. The average temperature, sunshine duration and the year of observation had negative association with BPD outbreak in Southwest, Nigeria (r = −0.420, −0.364 and −0.018, and r^2^ = 0.265, 0.360 and 0.035, respectively) as shown in Figs 4, 6 and 9. It was however observed that there was no relationship between the locations of these cocoa farms (Fig 7) and the specific period (month) when the disease was observed (Fig 8) with the outbreak of black pod disease in Nigeria.

**Fig 2: Relationship between BPD Outbreak and climatic factors in Southwest, Nigeria**

**Fig 3: Black pod disease outbreak and rainfall (1991-1995)**

**Fig 4: BPD Outbreak and average temperature in Southwestern Nigeria (1991-1995)**

**Fig 5: BPD Outbreak and average relative humidity in Southwestern Nigeria (1991-1995)**

**Fig 6: Black pod disease Outbreak and sunshine duration (1991-1995)**

**Fig 7: Black pod disease occurrence and sample station location (1991-1995)**

**Fig 8: Black pod disease Outbreak and period (in Months) of observation (1991-1995)**

**Fig 9: Black pod disease occurrence and the years of BPD documentation (1991-1995)**

### The Climate pattern of Southwest, Nigeria and its effects on black pod disease development

The climate pattern for Southwest, Nigeria in the late 1900s (20^th^ Century) showed that there was recurrent and substantial amount of rainfall experienced in Ogun, Ondo, Osun and Oyo States from March through October each year from 1991 to 1995 (Table 3). These periods possibly served as an interlude for proliferation and spread of the pathogen leading to possible infection of predisposed cocoa plants judging from the black pod disease occurrence report given by the Cocoa Research Institute of Nigeria (CRIN) from 1985-2014 as shown in Fig 10. Also, the climatology of the 21^st^ Century suggests that Ondo State had the highest amount of rainfall with annual rainfall value of 1,317.1mm and 1,381.0mm respectively in the year 2005 and 2006. The preference for heavy rainfall took a different precedence in 2007 and 2008, with Osun State leading the group for heavy down pour. The values for these years were 1,421.7mm and 1,597.6mm respectively. Oyo State, Osun State, and Ondo State took the lead for the succeeding years down to 2016 (Table 3). This accounted for the trend of disease outbreak with regards to the availability of moisture which is a pertinent factor for the survival and proliferation of the pathogen also reflected in the disease report from CRIN (Fig 10).

**Table 3:**
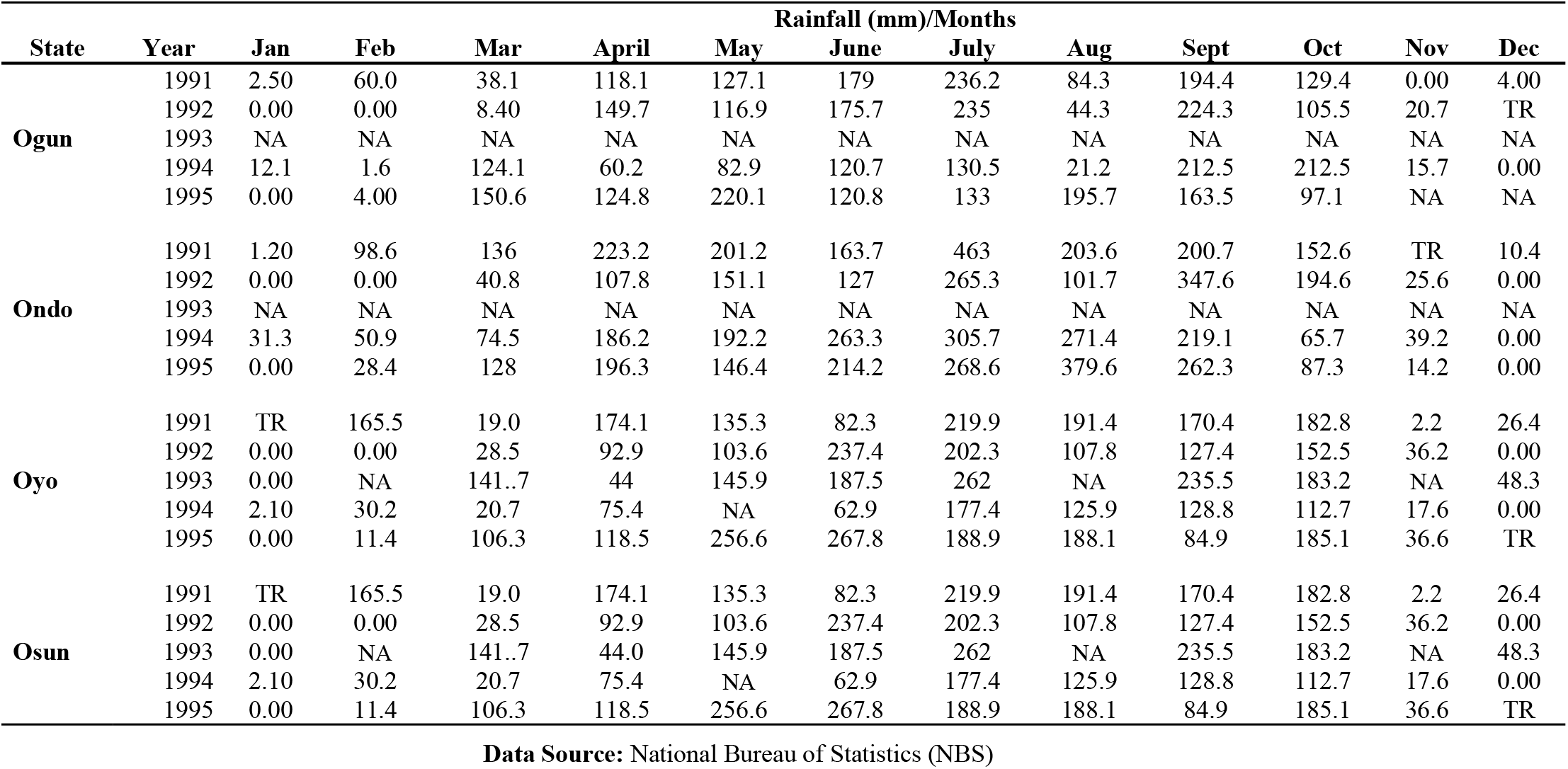
Monthly rainfall distribution for the Southwest, States of Nigeria

| State | Year | Rainfall (mm)/Months |  |  |  |  |  |  |  |  |  |  |  |
| --- | --- | --- | --- | --- | --- | --- | --- | --- | --- | --- | --- | --- | --- |
|  |  | Jan | Feb | Mar | April | May | June | July | Aug | Sept | Oct | Nov | Dec |
| Ogun | 1991 | 2.50 | 60.0 | 38.1 | 118.1 | 127.1 | 179 | 236.2 | 84.3 | 194.4 | 129.4 | 0.00 | 4.00 |
|  | 1992 | 0.00 | 0.00 | 8.40 | 149.7 | 116.9 | 175.7 | 235 | 44.3 | 224.3 | 105.5 | 20.7 | TR |
|  | 1993 | NA | NA | NA | NA | NA | NA | NA | NA | NA | NA | NA | NA |
|  | 1994 | 12.1 | 1.6 | 124.1 | 60.2 | 82.9 | 120.7 | 130.5 | 21.2 | 212.5 | 212.5 | 15.7 | 0.00 |
|  | 1995 | 0.00 | 4.00 | 150.6 | 124.8 | 220.1 | 120.8 | 133 | 195.7 | 163.5 | 97.1 | NA | NA |
| Ondo | 1991 | 1.20 | 98.6 | 136 | 223.2 | 201.2 | 163.7 | 463 | 203.6 | 200.7 | 152.6 | TR | 10.4 |
|  | 1992 | 0.00 | 0.00 | 40.8 | 107.8 | 151.1 | 127 | 265.3 | 101.7 | 347.6 | 194.6 | 25.6 | 0.00 |
|  | 1993 | NA | NA | NA | NA | NA | NA | NA | NA | NA | NA | NA | NA |
|  | 1994 | 31.3 | 50.9 | 74.5 | 186.2 | 192.2 | 263.3 | 305.7 | 271.4 | 219.1 | 65.7 | 39.2 | 0.00 |
|  | 1995 | 0.00 | 28.4 | 128 | 196.3 | 146.4 | 214.2 | 268.6 | 379.6 | 262.3 | 87.3 | 14.2 | 0.00 |
| Oyo | 1991 | TR | 165.5 | 19.0 | 174.1 | 135.3 | 82.3 | 219.9 | 191.4 | 170.4 | 182.8 | 2.2 | 26.4 |
|  | 1992 | 0.00 | 0.00 | 28.5 | 92.9 | 103.6 | 237.4 | 202.3 | 107.8 | 127.4 | 152.5 | 36.2 | 0.00 |
|  | 1993 | 0.00 | NA | 141..7 | 44 | 145.9 | 187.5 | 262 | NA | 235.5 | 183.2 | NA | 48.3 |
|  | 1994 | 2.10 | 30.2 | 20.7 | 75.4 | NA | 62.9 | 177.4 | 125.9 | 128.8 | 112.7 | 17.6 | 0.00 |
|  | 1995 | 0.00 | 11.4 | 106.3 | 118.5 | 256.6 | 267.8 | 188.9 | 188.1 | 84.9 | 185.1 | 36.6 | TR |
| Osun | 1991 | TR | 165.5 | 19.0 | 174.1 | 135.3 | 82.3 | 219.9 | 191.4 | 170.4 | 182.8 | 2.2 | 26.4 |
|  | 1992 | 0.00 | 0.00 | 28.5 | 92.9 | 103.6 | 237.4 | 202.3 | 107.8 | 127.4 | 152.5 | 36.2 | 0.00 |
|  | 1993 | 0.00 | NA | 141..7 | 44.0 | 145.9 | 187.5 | 262 | NA | 235.5 | 183.2 | NA | 48.3 |
|  | 1994 | 2.10 | 30.2 | 20.7 | 75.4 | NA | 62.9 | 177.4 | 125.9 | 128.8 | 112.7 | 17.6 | 0.00 |
|  | 1995 | 0.00 | 11.4 | 106.3 | 118.5 | 256.6 | 267.8 | 188.9 | 188.1 | 84.9 | 185.1 | 36.6 | TR |

Observations made from the prior studies conducted in the Plant Pathology/Mycology Laboratory of the Department of Botany, University of Ibadan, showed that *Phytophthora megakarya* thrived better when the ambient temperature ranged from 20°C to 30°C. The periods of the year that had maximum ambient temperatures corresponding to the optimum temperature requirement noted to support the metabolic activities of the pathogen were June, July, August, and September from 1991 to 1995 (Table 4). Sadly enough, these periods served as the peak periods for cocoa production in Southwest, Nigeria. Also, the minimum temperature all year round i.e. 1985 to 2013 (Fig 10) favoured the proliferation of the pathogen where other pertinent factors are available for prepenetration, penetration and infection (Table 5).

**Table 4:** Mean monthly maximum temperature reading for the Southwest, States of Nigeria

| State | Year | Max. Temperature (°C)/Months |  |  |  |  |  |  |  |  |  |  |  |
| --- | --- | --- | --- | --- | --- | --- | --- | --- | --- | --- | --- | --- | --- |
|  |  | Jan | Feb | Mar | April | May | June | July | Aug | Sept | Oct | Nov | Dec |
| Ogun | 1991 | 34.3 | 37.4 | 35.6 | 33.7 | 32.6 | 31.6 | 29.5 | 28.4 | 30.1 | 30.5 | 33.6 | 33.6 |
|  | 1992 | 34.5 | 37.3 | 36.3 | 35 | 32.7 | 30.3 | 38.3 | 28 | 29.2 | 31.9 | 33.2 | 34.8 |
|  | 1993 | 35.1 | 35.8 | 34.6 | 35 | 33.1 | 31.1 | NA | 29.7 | 30.8 | 32 | 33.6 | 33.8 |
|  | 1994 | 34 | 36.3 | 35.6 | 34.3 | 32.5 | 31.7 | 28.5 | 29.3 | 30.4 | 31.7 | 33.9 | 35.2 |
|  | 1995 | 35.5 | NA | NA | NA | 33.1 | 31.2 | NA | NA | NA | 31.5 | NA | NA |
| Ondo | 1991 | 32.2 | 33.4 | 32.6 | 31.3 | 30.7 | 29.6 | 28.2 | 26.9 | 28.8 | 29.3 | 31.8 | 32.1 |
|  | 1992 | 33 | 36 | 33.8 | 32.5 | 31.1 | 29 | 27 | 26.9 | 27.9 | 30 | 31.2 | 23.8 |
|  | 1993 | NA | NA | NA | NA | NA | NA | NA | NA | NA | NA | NA | NA |
|  | 1994 | 32 | 34.1 | 33.9 | 32.5 | 26.8 | 29 | NA | NA | 29 | 30.4 | 32.7 |  |
|  | 1995 | NA | NA | NA | NA | NA | NA | NA | NA | NA | NA | NA | NA |
| Oyo | 1991 | 33.5 | 34.9 | 34.6 | 33 | 31.6 | 31 | 29.3 | 27.7 | 28.1 | 30 | 32.2 | 32 |
|  | 1992 | 32.9 | 36.2 | 35.5 | 33.7 | 31.8 | 29.9 | 28 | 27.2 | 28.3 | 30.9 | 32.1 | 33.3 |
|  | 1993 | 33.1 | 34.6 | 33.5 | 33.1 | 32 | 30 | NA | 28.1 | 29.7 | NA | 31.9 | NA |
|  | 1994 | 32.7 | 34.9 | 35.5 | 34 | 32 | 30.7 | 27.9 | NA | 30 | 30.7 | 33.2 | 33.8 |
|  | 1995 | NA | NA | NA | NA | NA | NA | NA | NA | NA | NA | NA | NA |
| Osun | 1991 | 33.5 | 34.9 | 34.6 | 33 | 31.6 | 31 | 29.3 | 27.7 | 28.1 | 30 | 32.2 | 32 |
|  | 1992 | 32.9 | 36.2 | 35.5 | 33.7 | 31.8 | 29.9 | 28 | 27.2 | 28.3 | 30.9 | 32.1 | 33.3 |
|  | 1993 | 33.1 | 34.6 | 33.5 | 33.1 | 32 | 30 | NA | 28.1 | 29.7 | NA | 31.9 | NA |
|  | 1994 | 32.7 | 34.9 | 35.5 | 34 | 32 | 30.7 | 27.9 | NA | 30 | 30.7 | 33.2 | 33.8 |
|  | 1995 | NA | NA | NA | NA | NA | NA | NA | NA | NA | NA | NA | NA |

**Table 5:** Mean monthly minimum temperature reading for the Southwest, States of Nigeria

| State | Year | Minimum Temperature (°C)/Months |  |  |  |  |  |  |  |  |  |  |  |
| --- | --- | --- | --- | --- | --- | --- | --- | --- | --- | --- | --- | --- | --- |
|  |  | Jan | Feb | Mar | April | May | June | July | Aug | Sept | Oct | Nov | Dec |
| Ogun | 1991 | 23.8 | 26 | 25.2 | 23.7 | 24.2 | 23.8 | 23.1 | 22.7 | 22.8 | 22.6 | 24.2 | 22.5 |
|  | 1992 | 20.5 | 24.1 | 25.5 | 23.3 | 24 | 22.9 | 22.9 | 22.6 | 22.4 | 23.2 | 22.3 | 23.2 |
|  | 1993 | 21.1 | 24.5 | 23.7 | 24.5 | 24.2 | 23.5 | NA | NA | 22.8 | 23.3 | 23.8 | 22.2 |
|  | 1994 | 23.1 | 25.1 | 24.8 | 25.1 | 23.7 | 31.2 | 22.9 | 23 | 23.2 | 22.9 | 22.5 | 20.2 |
|  | 1995 | 22.2 | NA | NA | NA | 23.9 | 23.3 | NA | NA | NA | 23.3 | NA | NA |
| Ondo | 1991 | 19.6 | 22.6 | 22.7 | 21.2 | 21.9 | 21.2 | 21 | 21.3 | 21 | 20.2 | 21.1 | 18.2 |
|  | 1992 | 15.3 | 18.8 | 22.8 | 22.9 | 21.8 | 20.7 | 20.1 | 20.8 | 20.9 | 21.4 | 20.2 | 19 |
|  | 1993 | 17.3 | 20.6 | 21.9 | 23.1 | 22.9 | 22 | NA | 21.7 | 22.1 | NA | 22.1 | NA |
|  | 1994 | NA | NA | NA | NA | NA | NA | NA | NA | NA | NA | NA | NA |
|  | 1995 | 19.6 | 21.7 | 22.9 | 22.4 | NA | 21.2 | NA | NA | 21.7 | 21.2 | 20.7 | NA |
| Oyo | 1991 | 22.9 | 24.0 | 24.4 | 23.2 | 23.3 | 23 | 22.5 | 21.8 | 22.0 | 21.5 | 23.3 | 21.8 |
|  | 1992 | 20.2 | 22.9 | 24.3 | 23.8 | 23.3 | 22.9 | 22 | 21.4 | 21.7 | 22.1 | 21.9 | 22.4 |
|  | 1993 | 20.9 | NA | 22.9 | 23.5 | 23.3 | 22.4 | 22 | NA | 21.8 | 22.3 | NA | 22.2 |
|  | 1994 | 22.4 | 24.1 | 24.3 | 23.9 | 22.7 | 22.3 | 21.9 | NA | 22.6 | 22.2 | 22.4 | 20.4 |
|  | 1995 | NA | NA | NA | NA | NA | NA | NA | NA | NA | NA | NA | NA |
| Osun | 1991 | 22.9 | 24.0 | 24.4 | 23.2 | 23.3 | 23 | 22.5 | 21.8 | 22.0 | 21.5 | 23.3 | 21.8 |
|  | 1992 | 20.2 | 22.9 | 24.3 | 23.8 | 23.3 | 22.9 | 22 | 21.4 | 21.7 | 22.1 | 21.9 | 22.4 |
|  | 1993 | 20.9 | NA | 22.9 | 23.5 | 23.3 | 22.4 | 22 | NA | 21.8 | 22.3 | NA | 22.2 |
|  | 1994 | 22.4 | 24.1 | 24.3 | 23.9 | 22.7 | 22.3 | 21.9 | NA | 22.6 | 22.2 | 22.4 | 20.4 |
|  | 1995 | NA | NA | NA | NA | NA | NA | NA | NA | NA | NA | NA | NA |

It was observed form the climate pattern of the Southwest that from March through October from the early morning readings taken in 1991 to 1995 that a relatively humid atmosphere of 75% and above was recorded monthly which favoured the establishment of black pod disease suggesting the possibility of infection within these periods (Table 6-7). The periods of the year from 1991 to 1995 with afternoon Relative humidity readings of 75% and above was June, July, August and September across all the years investigated and within all the States analysed, favours disease proliferation (Fig 10). Also, the mean amount of saturated vapour present in the atmosphere from 1985 through 2014 also favoured the establishment of black pod disease in Southwest, Nigeria (Fig 10). The overall diagnosis was an indication of the parameter pertinent for disease establishment and when they combine favourably in favour of the noxious pathogen (*Phytophthora megakarya*), they can aid the proliferation, ramification and destruction of cocoa pods both ripe and unripe that falls within their path of travel.

**Table 6:** Relative Humidity values for the Southwest, States of Nigeria

| State | Year | Relative Humidity in the morning at 9.00GMT (%) / Months |  |  |  |  |  |  |  |  |  |  |  |
| --- | --- | --- | --- | --- | --- | --- | --- | --- | --- | --- | --- | --- | --- |
|  |  | Jan | Feb | Mar | April | May | June | July | Aug | Sept | Oct | Nov | Dec |
| Ogun | 1991 | 81 | 64 | 81 | 84 | 83 | 87 | 90 | 89 | 87 | 87 | 84 | 78 |
|  | 1992 | 55 | 70 | 76 | 81 | 83 | 85 | 89 | 89 | 88 | 86 | 76 | 78 |
|  | 1993 | 53 | 80 | 77 | 78 | 82 | 85 | NA | 87 | 87 | 85 | 89 | 78 |
|  | 1994 | 73 | 78 | 80 | 79 | 83 | 84 | 88 | 87 | 86 | 85 | 80 | 64 |
|  | 1995 | 87 | NA | NA | NA | 82 | 85 | NA | NA | NA | NA | 85 | NA |
| Ondo | 1991 | 75 | 79 | 81 | 83 | 84 | 85 | 89 | 89 | 86 | 84 | 78 | 67 |
|  | 1992 | 48 | 54 | 75 | 79 | 82 | 84 | 88 | 87 | 89 | 83 | 72 | 69 |
|  | 1993 | 48 | 68 | 71 | 76 | 77 | 82 | NA | NA | NA | NA | NA | NA |
|  | 1994 | 69 | 72 | 76 | 79 | NA | NA | NA | 85 | 85 | 82 | 71 | NA |
|  | 1995 | NA | NA | NA | NA | NA | NA | NA | NA | NA | NA | NA | NA |
| Oyo | 1991 | 70 | 78 | 76 | 81 | 81 | 83 | 88 | 88 | 85 | 84 | 79 | 70 |
|  | 1992 | 50 | 63 | 73 | 78 | 80 | 84 | 88 | 87 | 87 | 82 | 73 | 75 |
|  | 1993 | 48 | NA | 75 | 78 | 80 | 83 | 86 | NA | 85 | 83 | NA | 73 |
|  | 1994 | 68 | 73 | 74 | 75 | 82 | 81 | 89 | NA | 86 | 83 | 74 | 57 |
|  | 1995 | NA | NA | NA | NA | NA | NA | NA | NA | NA | NA | NA | NA |
| Osun | 1991 | 70 | 78 | 76 | 81 | 81 | 83 | 88 | 88 | 85 | 84 | 79 | 70 |
|  | 1992 | 50 | 63 | 73 | 78 | 80 | 84 | 88 | 87 | 87 | 82 | 73 | 75 |
|  | 1993 | 48 | NA | 75 | 78 | 80 | 83 | 86 | NA | 85 | 83 | NA | 73 |
|  | 1994 | 68 | 73 | 74 | 75 | 82 | 81 | 89 | NA | 86 | 83 | 74 | 57 |
|  | 1995 | NA | NA | NA | NA | NA | NA | NA | NA | NA | NA | NA | NA |

**Table 7:** Relative Humidity values for the Southwest of Nigeria

| Relative Humidity in the afternoon at 15.00GMT |  |  |  |  |  |  |  |  |  |  |  |  |  |
| --- | --- | --- | --- | --- | --- | --- | --- | --- | --- | --- | --- | --- | --- |
| State | Year | Jan | Feb | Mar | April | May | June | July | Aug | Sept | Oct | Nov | Dec |
| Ogun | 1991 | 48 | 54 | 38 | 63 | 72 | 73 | 79 | 80 | 74 | 73 | 35 | 47 |
|  | 1992 | 31 | 31 | 47 | 57 | 70 | 74 | 82 | 80 | 75 | 69 | 56 | 45 |
|  | 1993 | 28 | 41 | 50 | 58 | 67 | 75 |  | 74 | 75 | 67 | 59 | 48 |
|  | 1994 | 46 | 37 | 52 | 59 | 68 | 71 | 81 | 73 | 74 | 67 | 52 | 35 |
|  | 1995 | 35 | NA | NA | NA | 69 | 75 | NA | NA | NA | NA | 70 | NA |
| Ondo | 1991 | 44 | 50 | 55 | 64 | 71 | 71 | 79 | 81 | 73 | 70 | 53 | 40 |
|  | 1992 | 27 | 21 | 45 | 60 | 64 | 74 | 81 | 77 | 78 | 67 | 52 | 41 |
|  | 1993 | 28 | 35 | 48 | 55 | 64 | 66 | NA | NA | NA | NA | NA | NA |
|  | 1994 | 45 | 37 | 50 | 58 | NA | NA | NA | 75 | 75 | 69 | 53 | NA |
|  | 1995 | NA | NA | NA | NA | NA | NA | NA | NA | NA | NA | NA | NA |
| Oyo | 1991 | 43 | 49 | 50 | 58 | 67 | 69 | 78 | 79 | 71 | 67 | 54 | 45 |
|  | 1992 | 32 | 27 | 44 | 55 | 65 | 71 | 77 | 77 | 73 | 63 | 55 | 46 |
|  | 1993 | NA | NA | NA | NA | NA | NA | NA | NA | NA | NA | NA | NA |
|  | 1994 | 46 | 38 | 46 | 57 | 64 | 67 | 80 | NA | 72 | 66 | 49 | 36 |
|  | 1995 | NA | NA | NA | NA | NA | NA | NA | NA | NA | NA | NA | NA |
| Osun | 1991 | 43 | 49 | 50 | 58 | 67 | 69 | 78 | 79 | 71 | 67 | 54 | 45 |
|  | 1992 | 32 | 27 | 44 | 55 | 65 | 71 | 77 | 77 | 73 | 63 | 55 | 46 |
|  | 1993 | NA | NA | NA | NA | NA | NA | NA | NA | NA | NA | NA | NA |
|  | 1994 | 46 | 38 | 46 | 57 | 64 | 67 | 80 | NA | 72 | 66 | 49 | 36 |
|  | 1995 | NA | NA | NA | NA | NA | NA | NA | NA | NA | NA | NA | NA |

**Fig 10: Black pod disease pestilence in the Southwest of Nigeria**

### Development of prediction models for black pod disease in Nigeria

Several models were developed to assess the level of black pod disease development and its spread within the Southwest of Nigeria.

### Model 1 (MRM_1_)

General Equation (1991-1995)

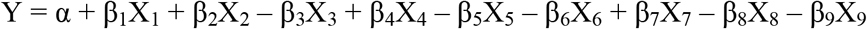

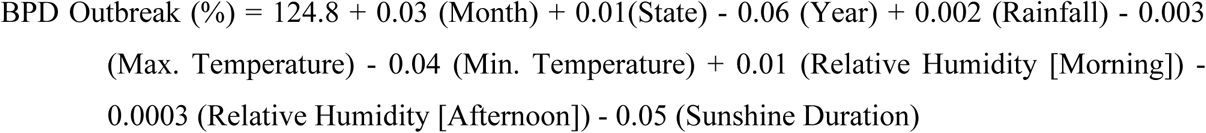

### Model 2 (MRM_2_)

General Equation (1991-1995)

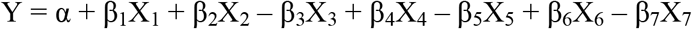

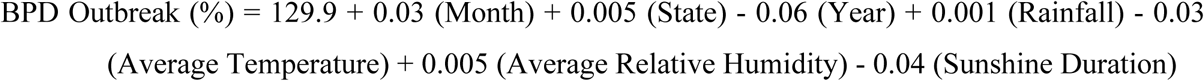

### Model 3 (MRM_3_)

General Equation (1991-1995)

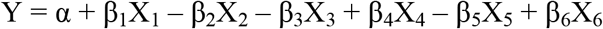

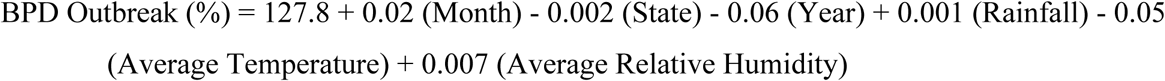

### Model 4 (MRM_4_)

General Equation (1991-1995)

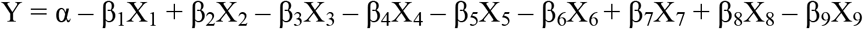

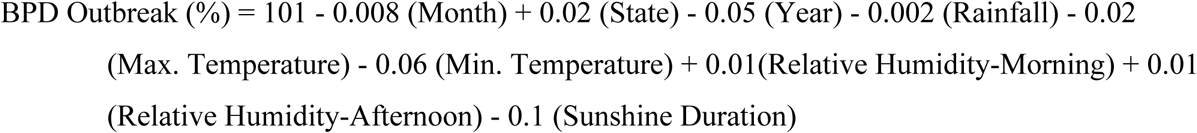

### Model 5 (MRM_5_) - ETAPOD

General Equation (1985-2014) [**Accepted Equation**]

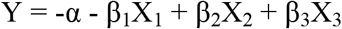

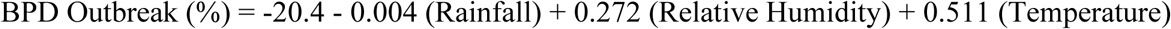

### Model 6 (MRM_6_)

General Equation (1991-1995)

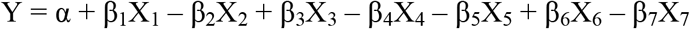

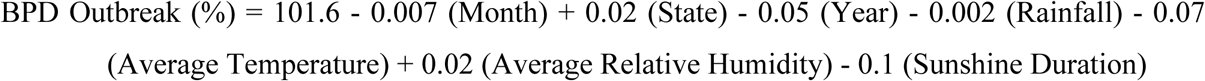

### Model 7 (MRM_7_)

General Equation (1985-2014)

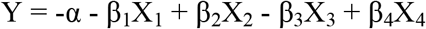

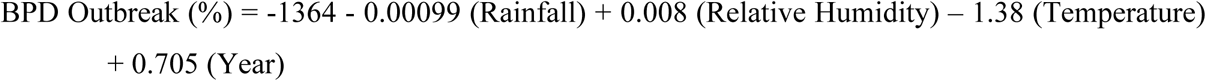

### Model 8 (MRM_8_)

General Equation (1991-1995)

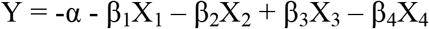

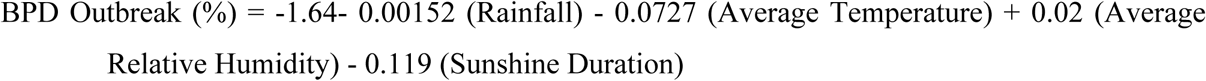

### Model Selection

Preliminary screening of the developed models was done using the co-efficient of correlation (R-Sq). The five (5) best fitted models (MRM_1_, MRM_2_, MRM_3_, MRM_4_, and MRM_5_) for black pod disease prediction were considered for further validation prior to final selection. The posthoc analysis conducted showed that MRM_5_ was the preferred model for black pod disease prediction followed by MRM_4_>MRM_1_>MRM_2_>MRM_3_ in terms of the Standard Error of Regression (SER) which was given as 0.22, 0.39, 0.45, 0.45, and 0.45 respectively; Root Mean Square Error of Prediction (RMSE_pred_): 0.30, 0.39, 0.46, 0.46 and 0.46 respectively; and the Adjusted Co-efficient of Correlation (R-Sq_Adj_): 0.67, 0.49, 0.32, 0.32 and 0.31 for MRM_5_, MRM_4_, MRM_1_, MRM_2_, and MRM_3_. The preferred model MRM_5_ was named “ETAPOD” (Fig 11)

**Fig 11: MRM_5_ BPD prediction Model (ETAPOD)**

### Prediction of black pod disease and validation of results

The predicted level of black pod disease outbreak for Ogun State was 9.97% in May, June (11.54%), July (12.25%), August (11.24%), September (9.86%), October (9.24%), November (5.95%), December (2.25%) in 2015 and 1.03% for January, February (2.81%), March (4.74%), April (7.42%), May (9.97%), in 2016 (Fig 12). That of Ondo State was predicted thus: May, 2015 (8.58%); June, 2015 (9.05%); July, 2015 (11.48%); August, 2015 (10.26%); September, 2015 (10.09%); October, 2015 (8.17%); November, 2015 (4.50%) and December, 2015 (0.76%). While the predictions for 2016 was given thus: January (−1.40%), February (−0.04%), March (4.32%), April (6.48%), and May (8.58%), respectively (Fig 13).

For Osun State, black pod disease outbreak was predicted in 2015 as follows: May (8.64%), June (9.43%), July (11.82%), August (10.34%), September (10.26%), October (7.80%), November (4.94%), and December (1.67%); that of 2016 was predicted thus January (0.04%), February (1.25%), March (4.69%), April (6.89%) and May (8.64%) as shown in Fig 14. Finally, the predictions for Oyo State was as follow: May (8.69%), June (9.43%), July (11.77%), August (10.39%), September (9.98%), October (7.80%), November (4.95%), December (1.67%) for 2015 and January (0.21%), February (1.29%), March (4.57%), April (6.87%) and May (8.69%) for 2016 growing season (Fig 15). A comparison was drawn with the observed values obtained in the field for the 2015/2016 cocoa production season.

### Result Validation: Predicted (Computer Simulations) versus Observed BPD Outbreak

The predicted and actual black pod disease occurrence within the States where the study areas where located were compared to determine their level of accuracy. The major season for cocoa production was considered. Black pod disease outbreak for Ondo in the month of June was predicted as 9.05% and the actual observation made in the field was 9.5%, it was predicted as 11.5% in July (Actual observation was 18.0%), in August, predicted result of BPD outbreak was 10.3% (Actual BPD outbreak was 26.5%), in September (Predicted BPD Outbreak = 10.1%, Actual BPD Outbreak =11.0%), and in October (Predicted BPD Outbreak = 8.17%, while Actual BPD Outbreak = 5.0%) as stated in Table 8.

**Table 8:**
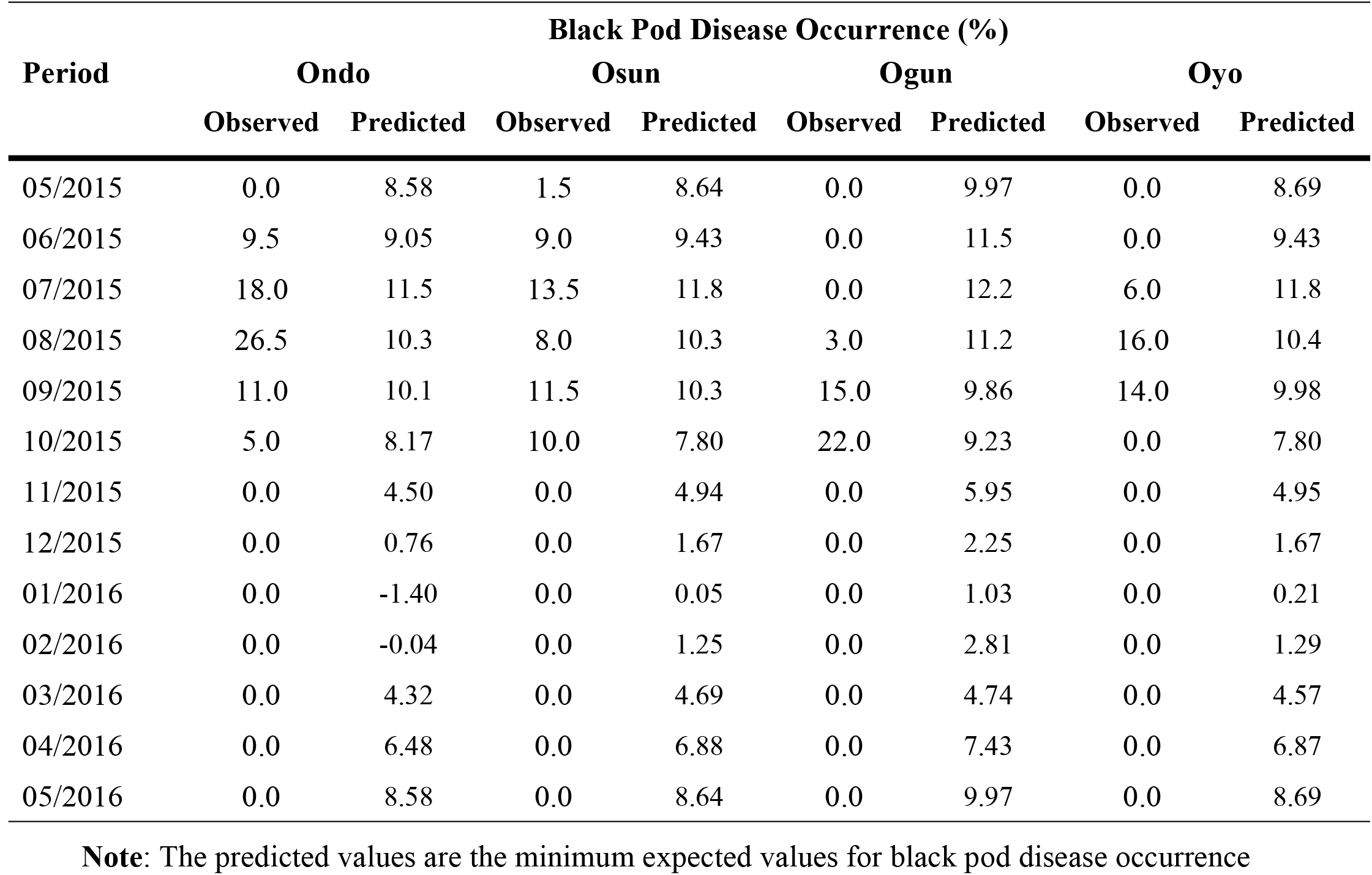
BPD outbreak both in the field and feedback information from MRM_5_ (ETAPOD)

In Osun, the predicted BPD Outbreak for June was 9.43% (Actual BPD Outbreak = 9.0%), in July (Predicted BPD Occurrence = 11.8%, Actual BPD Occurrence = 13.5%), August (Predicted BPD Outbreak = 10.3%, Actual BPD Incidence = 8.0%), in September (Predicted Outbreak for black pod disease = 10.3%, Actual Value = 11.5%), and October (Predicted Result = 7.8%, Actual Occurrence = 10.0%). The predictions of black pod disease made by ETAPOD for Ogun was [June (Predicted BPD Incidence = 11.5%, Actual BPD Occurrence = 0.0%), July (Predicted BPD Incidence = 12.2%, Actual BPD Incidence = 0.0%), August (Predicted BPD Incidence = 11.2%, Actual BPD Outbreak = 3.0%), September (Predicted BPD Outbreak = 9.86%, Actual BPD Occurrence = 15.0%), and October (Predicted BPD Outbreak = 9.23%, Actual BPD Outbreak = 22.0%)]. Finally, that of Oyo State was given thus: June (9.43%, 0.0%), July (11.8%, 6.0%), August (10.4%, 16.0%), September (9.98%, 14.0%), and October (7.8%, 0.0%) for both predicted and actual black pod disease outbreak (Table 9). Predictions on BPD outbreak made by ETAPOD showed that the Monthly BPD Outbreak (MBO), the Total Annual BPD Outbreak (TAO) and the Average Annual BPD Outbreak (AAO) for some selected stations i.e. Ọwenà and Wáàsimi were 9.05, 72.3 and 6.0% compared with observed BPD values of 9.5, 70.0, and 5.8%, respectively. Adaàgbà, Iyánfọwọrọgi, and Owódé-Igàngán had 9.43, 77.8, and 6.5% as their predicted BPD values compared with the observed values of 9.0, 53.5, and 4.46%, respectively as shown in Figs 12-15.

**Table 9:** An estimation of the difference that exist between the data set

| Period | Estimated Difference (%) |  |  |  |
| --- | --- | --- | --- | --- |
|  | Ondo | Osun | Ogun | Oyo |
| 05/2015 | -8.58 | -7.14 | -9.97 | -8.69 |
| 06/2015 | 0.45 | -0.43 | -11.5 | -9.43 |
| 07/2015 | 6.50 | 1.70 | -12.2 | -5.80 |
| 08/2015 | 16.2 | -2.30 | -8.20 | 5.60 |
| 09/2015 | 0.90 | 1.20 | 5.14 | 4.02 |
| 10/2015 | -3.17 | 2.20 | 12.8 | -7.80 |
| 11/2015 | -4.50 | -4.94 | -5.95 | -4.95 |
| 12/2015 | -0.76 | -1.67 | -2.25 | -1.67 |
| 01/2016 | 1.40 | -0.05 | -1.03 | -0.21 |
| 02/2016 | 0.04 | -1.25 | -2.81 | -1.29 |
| 03/2016 | -4.32 | -4.69 | -4.74 | -4.57 |
| 04/2016 | -6.48 | -6.88 | -7.43 | -6.87 |
| 05/2016 | -8.58 | -8.64 | -9.97 | -8.69 |

**Plate 12: BPD outbreak predictions in Ogun State (2015/2016)**

**Plate 13: BPD outbreak predictions in Ondo State (2015/2016)**

**Plate 14: BPD outbreak predictions in Osun State (2015/2016)**

**Plate 15: BPD outbreak predictions in Oyo State (2015/2016)**

### Performance of MRM_5_ forecast model (ETAPOD)

It was also observed that the range of disparity between the observed and predicted values for Ondo State was between −8.58% and 16.2%, Osun (−7.14% and 2.20%), Ogun (−11.5% and 12.8%) and Oyo (−9.43% and 5.60%) as recorded in Table 9. The estimated performance of the developed black pod disease occurrence forecast model was rated as follows: Ondo State had good black pod disease predicted values for the months of June, July, August, September 2015, January and February 2016; whereas, fair black pod disease occurrence was experienced in the months of October, November, December 2015 and March, 2016. Osun State had good black pod disease predicted values for the months of July, September, October 2015, and January 2016; and fair black pod disease predicted values for the months of June, August, November, December 2015, February and March 2016. Ogun State had black pod disease values predicted correctly for the months of September and October 2015 only; whereas, there was a series of fair black pod incidence predicted values within the months of December 2015, January, February and March 2016. Finally, for Oyo State there was good black pod disease prevalence values predicted for the months of August and September 2015 only, and fair predicted black pod disease occurrence values for the months of November, December 2015, January, February, and March 2016 as shown in Table 10.

**Table 10:** Performance rating for MRM_5_ forecast model (ETAPOD)

| Quality of black pod disease occurrence prediction |  |  |  |  |
| --- | --- | --- | --- | --- |
| Period | Ondo | Osun | Ogun | Oyo |
| 05/2015 | - | - | - | - |
| 06/2015 | + | -/+ | - | - |
| 07/2015 | + | + | - | - |
| 08/2015 | + | -/+ | - | + |
| 09/2015 | + | + | + | + |
| 10/2015 | -/+ | + | + | - |
| 11/2015 | -/+ | -/+ | - | -/+ |
| 12/2015 | -/+ | -/+ | -/+ | -/+ |
| 01/2016 | + | + | -/+ | -/+ |
| 02/2016 | + | -/+ | -/+ | -/+ |
| 03/2016 | -/+ | -/+ | -/+ | -/+ |
| 04/2016 | - | - | - | - |
| 05/2016 | - | - | - | - |

### The statistical error of prediction for the developed black pod disease occurrence prediction model

The error of prediction was estimated statistically and the level of accuracy of the developed model for prediction of black pod disease occurrence was determined by simple statistical formula. The estimation of the monthly percentage error of prediction for each state was given thus: for Ondo State it was estimated to be 0.20% in the month of June 2015, July 2015 (42.25%), September 2015 (0.81%), October 2015 (10.05%), November 2015 (20.25%), December 2015 (0.58%), January 2016 (1.96%), February 2016 (0.0%), March 2016 (18.66%), and April 2016 (41.99%) as stated in Table 11.

**Table 11:** Percentage error in black pod disease prediction

| Periods | Error in prediction of black pod disease occurrence (%) |  |  |  |
| --- | --- | --- | --- | --- |
|  | [E= (Y-Ŷ)<sup>2</sup>] |  |  |  |
|  | Ondo | Osun | Ogun | Oyo |
| 05/2015 | 73.62 | 50.98 | 99.40 | 75.52 |
| 06/2015 | 0.20 | 0.18 | 132.25 | 88.92 |
| 07/2015 | 42.25 | 2.89 | 148.84 | 33.64 |
| 08/2015 | 262.44 | 5.29 | 67.24 | 31.36 |
| 09/2015 | 0.81 | 1.44 | 26.42 | 16.16 |
| 10/2015 | 10.05 | 4.84 | 163.84 | 60.84 |
| 11/2015 | 20.25 | 24.4 | 35.4 | 24.5 |
| 12/2015 | 0.58 | 2.79 | 5.06 | 2.79 |
| 01/2016 | 1.96 | 0.00 | 1.06 | 0.04 |
| 02/2016 | 0.00 | 1.56 | 7.90 | 1.66 |
| 03/2016 | 18.66 | 22.00 | 22.47 | 20.88 |
| 04/2016 | 41.99 | 47.33 | 55.2 | 47.2 |
| 05/2016 | 73.62 | 74.65 | 99.4 | 75.52 |

It was noted that the statistical error of prediction of black pod disease occurrence in Osun State was low with the value estimated for the month of June 2015 being 0.18%, July 2015 (2.89%), August 2015 (5.29%), September 2015 (1.44%), October 2015 (4.84 %), November 2015 (24.4%), December 2015 (2.79%), January 2016 (0.0%), February 2016 (1.56%), March 2016 (22.0%), and April 2016 (47.33%) during the 2015/2016 cocoa production season across the Southwest of Nigeria (Table 11).

Ogun and Oyo States had similar estimated statistical error in black pod disease prediction. Although their estimated levels in the error of black pod disease prevalence predicted values were low, much work still need to be done to improve the quality of the result forecasted for these states. It was noted that the error of prediction for Ogun State was 26.42% for the month of September 2015, 35.4% for November 2015, December 2015 (5.06%), January 2016 (1.06%), February 2016 (7.9%), and March 2016 (22.47%). The statistical error of prediction for Oyo State was 33.64% for the month of July 2015, August 2015 (31.36%), September 2015 (16.16%), November 2015 (24.5%), December 2015 (2.79%), January 2016 (0.04%), February 2016 (1.66%), March 2016 (20.88%), and April 2016 (47.2%) as estimated in the 2015/2016 cocoa production season across the Southwest, states of Nigeria (Table 11).

## Discussion

### Weather survey in line with BPD outbreak in Southwestern Nigeria

The weather report in the early 1900s for Southwestern Nigeria showed that there was recurrent rainfall within the months of March through October from 1991 to 1995. Also, ambient temperature was low during the day and at night, and there was much saturated water vapour in the air across the four (4) States investigated within the same period. March to October happen to be the most productive periods for Cocoa production in Southwest, Nigeria; Therefore, the observations noted gives an indication on the possibility of infection within these periods. This favourable weather pattern for black pod disease infection was earlier reported by Akrofi [8].

### The required predictor variables

The most pertinent factors needed for structuring a forecast model for the prediction of BPD outbreak in Southwest, Nigeria include weather reports for rainfall, temperature, relative humidity and sunshine duration spread across the years, a good source of cocoa yield across the selected region(s), and data recorded for black pod disease pestilence. This was in line with the requirements stipulated by Fernandes [9] as pertinent factors needed for the establishment of a good warning system.

### BPD forecast model structuring

The Multiple Regression Model (MRM): Y=α+β_1_X_1_+β_2_X_2_+…+β_n_X_n_ where Y is Nx1 matrix of response variable, X_1_,X_2_,…X_n_ are NxK matrices of regressors, and β_1_,β_2_, …β_n_ regression coefficients was used in model development. Eight models (MRM_1_-MRM_8_) were fitted from real life BPD data. The performances of the models were ascertained using SER, RMSE_pred_ and R-Sq_Adj_. This was as prescribed by Simon [10], Luo [11] and Wikipedia [7].

### Accuracy of BPD outbreak predictions by ETAPOD

The MRM_5_ model (ETAPOD) was able to predict BPD outbreak accurately in Ondo and Osun States for the major production season in Nigeria, but the predictions made for Ogun and Oyo States were slightly inaccurate suggesting and improvement in model quality. The research information generated from ETAPOD was in line with the observations made by Opoku *et al*. [1] and [12] in their research conducted in Ghana. He stated that primary infections of BPD usually occur around June, but the peak of infection generally occurs between August and October. Information on peak periods for BPD infection can be useful when planning management strategies for BPD eradication. Luo [11] also designed a forecast model for the prediction of foliar diseases of winter wheat caused by *Septoria tritici* across England and Wales and his predictions for the disease was seemingly not 100% accurate.

## Supporting Information

**S1 Fig: BPD outbreak and the presence of abundant inoculum of the pathogen**

**S2 Fig: Rainfall pattern and how it affects BPD outbreak**

**S3 Fig: The effect of increasing temperature and BPD outbreak**

**S4 Fig: An increase in saturated vapour in the air and how it influence BPD outbreak**

**S5 Fig: Sunshine duration and how it influences BPD proliferation**

**S6 Fig: Wind Speed and BPD outbreak**

**S7 Fig: Period of cocoa pod formation/maturation and BPD infection**

**S8 Fig: atmospheric pressure and BPD outbreak**

## Recommendation

ETAPOD harnesses several potentials and possibilities that can be improved on to obtain excellent results. The accuracy of the warning system developed for the prediction of black pod disease (ETAPOD) can be perfected if:

1. Weather parameters are obtained from meteorological stations situated in the farm or those closely located to the region where active cocoa production take place.
2. The level of accuracy of predicted weather reports is above 95%
3. Consistency of cocoa production within that locality is constant
4. The type of cropping system employed could be determined
5. cocoa is the major crop cultivated on the piece of land
6. Advanced digital image analysis could be used to improve measurement precision of disease prevalence and severity.

## Conclusion

ETAPOD harnesses the potentials to improve the functionality of other existing management strategies for the control of black pod disease in Nigeria by providing information regarding black pod disease occurrence, detect areas under severe attack by the disease (AUSA), and discourage fungicide misuse among local cocoa farmers. ETAPOD is unique in the sense that its primary functions in terms of black pod disease prediction are not geographically bound by location and as such the developed programme can be manipulated to provide optimum results anywhere needed in Nigeria, Africa and all around the world. Its ability to provide qualitative and quantitative description of the black pod disease pressure makes it superior to other forms of black pod disease control strategies in use.

Therefore, ETAPOD is a pertinent tool that can effectively minimize the prevalence of black pod disease of cocoa within Nigeria with minimal chemical application, decreasing the risk of chemical poisoning and increasing the production of healthy cocoa products nationwide. This is the surest and fastest way to ensuring sustainability of cocoa production in Nigeria and the world at large as a means to tackle the problem of food scarcity and unavailability of raw materials for production.

